# Sex-specific gut bacteria manipulation impacts offspring production in the honeybee parasite small hive beetle, *Aethina tumida* Murray

**DOI:** 10.1101/2023.10.15.562436

**Authors:** Md Jamil Biswas, Md Kawsar Khan, Sahareh Bashiribod, Fleur Ponton

## Abstract

Symbiotic bacteria play crucial roles in growth, development, behaviour, immunity, and metabolism of insects. Most of the research until now has explored the role of gut bacteria in insect model systems. The extent to which gut bacteria affect mating behaviour and reproduction of insect pest species still needs, however, to be fully comprehended. In this study, we aimed to investigate the effects of gut bacteria manipulation on mating behaviour and fecundity of a major beehive pest, the small hive beetle *Aethina tumida* (Murray). We manipulated gut bacteria of both sexes and determined the effects on mating behaviour and reproductive output. Our results provided evidence that manipulation of the gut bacteria influences reproduction in both sexes, negatively affecting mating latency, mating number, mating duration and offspring number (larvae). These results highlight the role of gut bacteria on the reproduction of insect pests, which might help in designing new pest control methods.

## 1 INTRODUCTION

Understanding the bacterial-mediated processes on insect reproduction is important because it can lead to novel control strategies for economically important insect pest species (Engl & Kaltenpoth, 2018). Diverse microbial communities affect the physiology and behaviour of insects (Engel & Moran, 2013; Jordan & Tomberlin, 2021; Lewis & Lizé, 2015; Rowe, Veerus, Trosvik, Buckling, & Pizzari, 2020). Among these, the gut bacteriome is crucial for a range of insect functions such as metabolism (Newell & Douglas, 2014; Wong, Dobson, & Douglas, 2014), development (Ridley, Wong, Westmiller, & Douglas, 2012; Shin et al., 2011; Storelli et al., 2011), nutrition (Douglas, 2009; Téfit & Leulier, 2017), immunity (Buchon, Broderick, Poidevin, Pradervand, & Lemaitre, 2009; Lee et al., 2013; Sansone et al., 2015), and longevity (Clark et al., 2015). In recent years, the role of symbiotic bacteria in insects has drawn increasing attention owing to their importance for conservation (Zhao, Lin, & Guo, 2022) and management (Crotti et al., 2012; Qadri, Short, Gast, Hernandez, & Wong, 2020). For example, volatiles from symbiotic bacteria are used as insect semiochemicals for monitoring and managing pest species (Davis, Crippen, Hofstetter, & Tomberlin, 2013; Leroy et al., 2011). The bacterial communities residing in the gut also play an important role in insect chemical communication through influencing the production of aggregation (Dillon et al. 2002, Wada-Katsumata et al. 2015) and sex pheromones (Hoyt, Osborne, & Mulcock, 1971; Sharon et al., 2010). For example, a *Bacillus* species isolated from male oriental fruit fly rectum, can produce male-borne sex pheromones when provided with adequate substrates (Ren, Ma, Xie, Lu, & Cheng, 2021). Furthermore, in *Drosophila melanogaster*, manipulation of the gut bacteria may impact insect mating preference (Sharon et al. 2010, 2011) and oviposition behaviour (Chaudhury, Skoda, Sagel, & Welch, 2010; Chaudhury, Zhu, & Skoda, 2016). The gut bacteria also influence mating duration and offspring production in female *Drosophila* with some strains negatively affecting male fertility when exclusively present in the gut (Morimoto, Simpson & Ponton, 2017).

The small hive beetle (SHB) *Aethina tumida* Murray (Coleoptera: Nitidulidae) is an opportunistic parasite that feeds on larvae, honey, and pollen of European honeybees, *Apis mellifera* L. (Hymenoptera: Apidae) (Murray, 1867). It has been reported that small hive beetle may also infect colonies of bumble bees (Hoffmann, Pettis, & Neumann, 2008) and stingless bees (Halcroft, Spooner-Hart, & Neumann, 2011). Small hive beetle is a worldwide honey bee pest (Giangaspero & Turno, 2015) and is now the predominant apiary pest in warm, moist regions of Eastern Australia (Leemon et al., 2018). Because this parasitic beetle mainly feed on hive material, they acquire bee symbiotic microbes that colonize their digestive tract (Huang, Lopez, & Evans, 2019). Twenty bee-associated bacteria have been detected in small hive beetle larvae and adults (Huang, Lopez, & Evans, 2019). Further 55 and 22 unique bacteria that are absent in bees were detected in small hive beetle larvae and adults respectively (Huang, Lopez, & Evans, 2019). Most of the microbes residing in small hive beetles are now known, however, how gut symbionts influence host traits such as reproduction is unclear.

In this study, we aim to determine the role of gut bacterial communities on mating behaviour and reproduction in small hive beetles. We manipulated gut bacteria in males and females and performed mating experiments where we measured mating latency, mating number, mating duration, and offspring number. Our results provided evidence that gut bacteria impact small hive beetle behaviour and reproduction. Our results are discussed in the context of applying these results to develop new control methods for this major honey bee pest.

## 2 MATERIALS AND METHODS

### 2.1 Insects

Small hive beetles were collected as adults from beehives at Macquarie University, New South Wales, Australia. Adult beetles were reared in laboratory colonies for seven generations and maintained in a controlled environment room at 26 ± 0.5 °C with 65 ± 5% relative humidity and 12 hours dark and 12 hours light photoperiod in plastic boxes (11 cm x 31 cm x 22 cm). Beetles were provided an artificial diet made with sugar, bee pollen (Hornsby Beekeeping Supplies, Australia), and pollen substitute (BeeSmart Beekeeping, Australia) in a 1:1:2:1 volume ratio with ad libitum water (Neumann et al., 2013). The wandering larvae were transferred into glass jars (21.1 cm x 16 cm) containing autoclaved soil and sand (1:1) for pupation (Neumann et al., 2013).

### 2.2 Bacterial manipulation

Beetles were allocated to three treatments: control group (intact gut bacteria), axenic group (very low number of gut bacteria), and re-inoculation group (axenic line reinoculated with the microbiota of control beetles). Beetle eggs were collected from generation seven in the lab-adapted colony, then dechorionated for 3 min in 0.5% bleach (Peerless JAL®), followed by one wash in 70% ethanol for 1 min and three washes in sterile Milli-Q water for generating axenic group. The control group was treated by washing eggs in sterile Milli-Q water three times. Control eggs (0.5 mL) were crushed in 1 mL sterile PBS in a sterile centrifuge tube using a hand-holding pestle cordless motor (Sigma, cat no. Z359971) and inoculated to axenic eggs for generating reinoculated lines. A sterile diet was prepared by autoclaving the standard diet. Eggs were then spread on the sterile diet in a glass box (2.7 L) (three replicates for each treatment group). Eggs were reared to the larval stage in a PCR cabinet (AirClean® 600 PCR Workstation). Third instar larvae were transferred into glass jars containing sterile media (50% soil + 50% sand) and kept in the PCR cabinet for pupation. Beetles were sexed one day after emergence, and males and females were kept separately in mesh bugDorm cages (32.5 × 32.5 × 32.5 cm). The bacterial status of eggs, larvae, and adult beetles was assessed by counting bacterial colonies on Luria Bertani agar (LB, Life technologies, cat no. 22700–025) media plates. To determine the bacterial status of adults, three adult male and female beetles (2 days after emergence), from each treatment group, were collected separately in two sterile microcentrifuge tubes. The beetles’ bodies were then washed once in 70% ethanol (Sigma, cat no. 64175) for 1 min, followed by three washes in sterile water. Afterward, 750 μl PBS buffer was added to each tube, and beetles were crushed using hand-holding pestle cordless motor (Sigma, cat no. Z359971). Both tubes were centrifuged (Benchmark MyFuge™ Mini) for a minute (6000 rpm) to separate the supernatant from the rest of the body debris. The supernatant was transferred to a centrifuge tube (1 mL). The solution was then diluted into two concentrations (10^−1^ and 10^−2^). Each suspension was pipetted (20 μl) and spread using a sterile spreader on agar plate media (LB) using single-use L-shape spreaders (Sigma, cat no. Z723193) and cultured at 30°C for 48 h. To determine the bacterial status of the eggs and larvae, fifty eggs and one larva were collected from each treatment group for each replicate. The same method as for the adult beetles was then followed. After counting the bacterial colonies on LB agar plates, we confirmed that the axenic adult group (both male and female) exhibited a very low number of bacterial colonies across all three replicates (maximum of n = 12 colonies from 20 μl of suspension on one agar plate media) than the control and reinoculated group (maximum of n = 58 colonies from 20 μl of suspension on one agar plate media) for each concentration (10^−1^ and 10^−2^). No bacteria was observed at the egg stage of the axenic group whereas a maximum of n = 2 colonies from 20 μl of suspension on one agar plate media were found in the larval stage of the axenic group. A maximum of n= 35 colonies from 20 μl of suspension on one agar plate media were observed in the control and reinoculated larvae.

### 2.3 Mating behaviour, and offspring production

Mating behaviour (mating latency, mating number, and mating duration) was measured for 10-day-old beetles. Mating experiments were performed for nine crossing groups that included three treatments (control, axenic, and reinoculated) for males and females. Twenty replicates were run in each crossing group. Pairs of adult beetles were transferred into Petri dishes (90-mm petri dish, cat. no., S6014S10, Techno Plas) containing ∼10 g of autoclaved artificial diet and mating behaviour was observed for fifteen minutes. After mating experiment, males were removed, and females were kept in the Petri dishes to lay egg. Beetle pairs that did not mate within 15 minutes were not kept for the egg laying experiment. Two microscope slides with a coverslip were kept inside each petri dish, providing a surface for egg-laying (Neumann et al., 2013). Based on our previous observations, we noticed that some females laid eggs three to four days after mating, so mated females were kept for 5 days to examine whether they laid eggs. Mated females that laid eggs in the 5 days were included in the analysis for the larvae count. Eggs were kept in Petri dishes until larvae hatched and the number of first instar larvae was recorded.

### 2.5 Statistical analysis

We performed all analyses in R (v.4.2.2) (R Core Team, 2019). All tests were run with sex and treatment as explanatory variables. We applied a generalized linear model (GLM) (zero-inflated negative binomial family to account for overdispersion) to analyse larvae number, a GLM (binomial family) to analyse mating occurrence (yes/no), a GLM (quasipoisson family to account for overdispersion) to analyse the total number of matings, a GLM (gaussian family) for the average latency period for each couple, and the mating duration (boxcox transformed).

## 3 RESULTS

### 3.1 Larvae number

The interaction between male and female treatment influenced the number of larvae (GLM (zero-inflated negative binomial): Male: χ^2^ = 7.184, *df* = 2, *p* = 0.027, Female: χ2 = 5.345, *df* = 2, *p* = 0.069, Male*Female: χ2 = 15.219, *df* = 4, *p* < 0.01). Axenic females produced a low number of larvae, whatever the treatment of their male partner (Figure 1). Control females had fewer larvae when mated with axenic males than those mated with control and reinoculated males (Figure 1). Reinoculated females showed a greater number of larvae when mated with control males and a slightly lower number of larvae when coupled with either axenic or reinoculated males (Figure 1).

**Figure 1.**
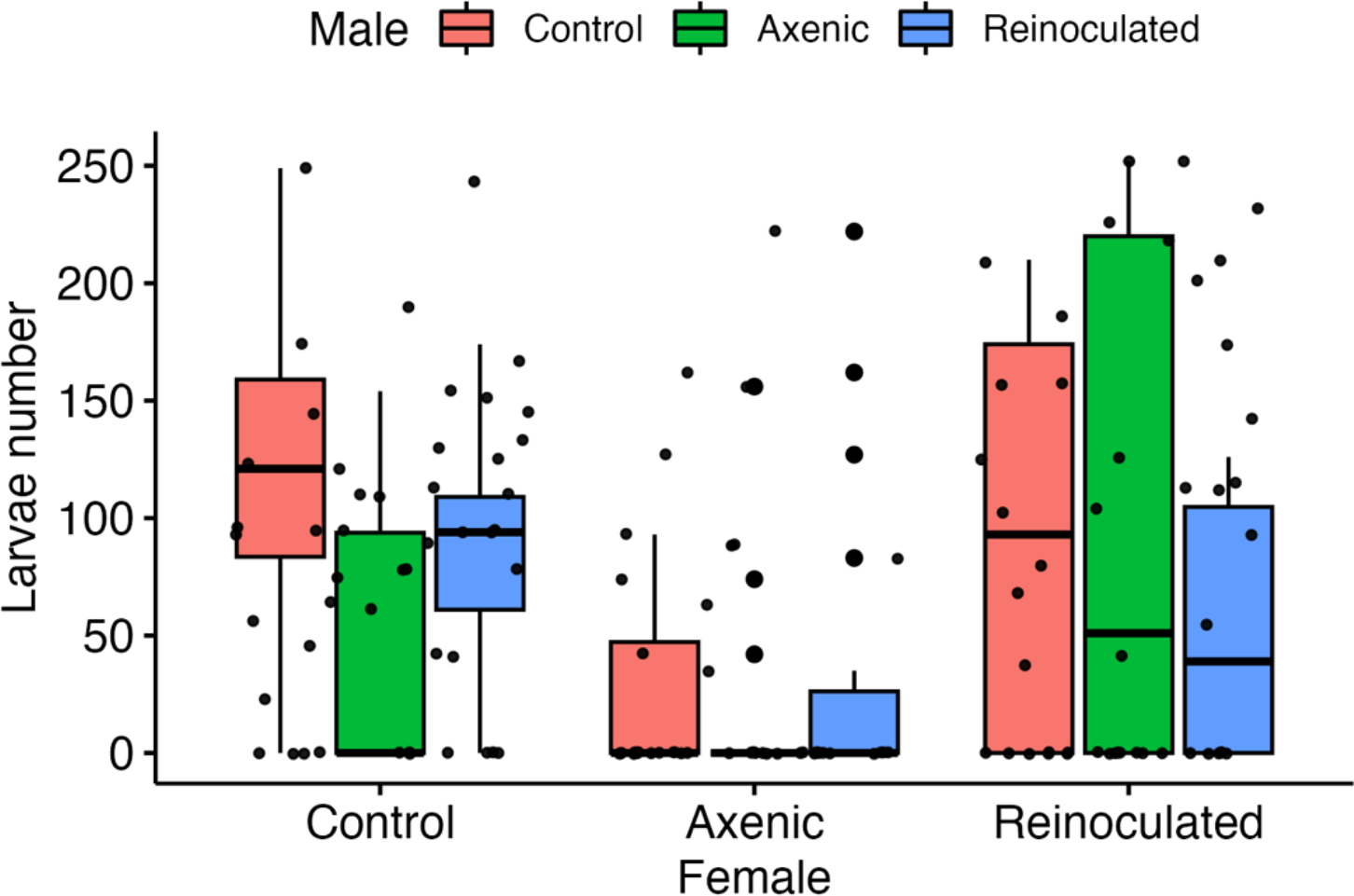
Number of larvae produced by pairs between control, axenic, and reinoculated female and male hive beetles. Boxplots depict the median, the 25^th^ and 75^th^ percentile with the whiskers extended to the minimum and maximum data points.

### 3.2 Mating occurrence

Bacterial manipulation in male or female did not influence mating occurrence, which means that all couples intended mating (GLM (binomial): Male: res.dev = 177.506, *df* = 2, *p* = 0.267; Female: res.dev = 176.884, *df* = 2, *p* = 0.732, Male*Female = res.dev = 174.326, *df* = 4, *p* = 0.634).

### 3.3 Mating number

The interaction between male and female treatment influenced the total number of mating events for each couple (GLM (quasipoisson): Male: *F* = 2.257, *df* = 2, *p* = 0.108; Female: *F* = 1.847, *df* = 2, *p* = 0.161; Male*Female: *F* = 3.706, *df* = 4, *p* < 0.01; Figure 2).

**Figure 2.**
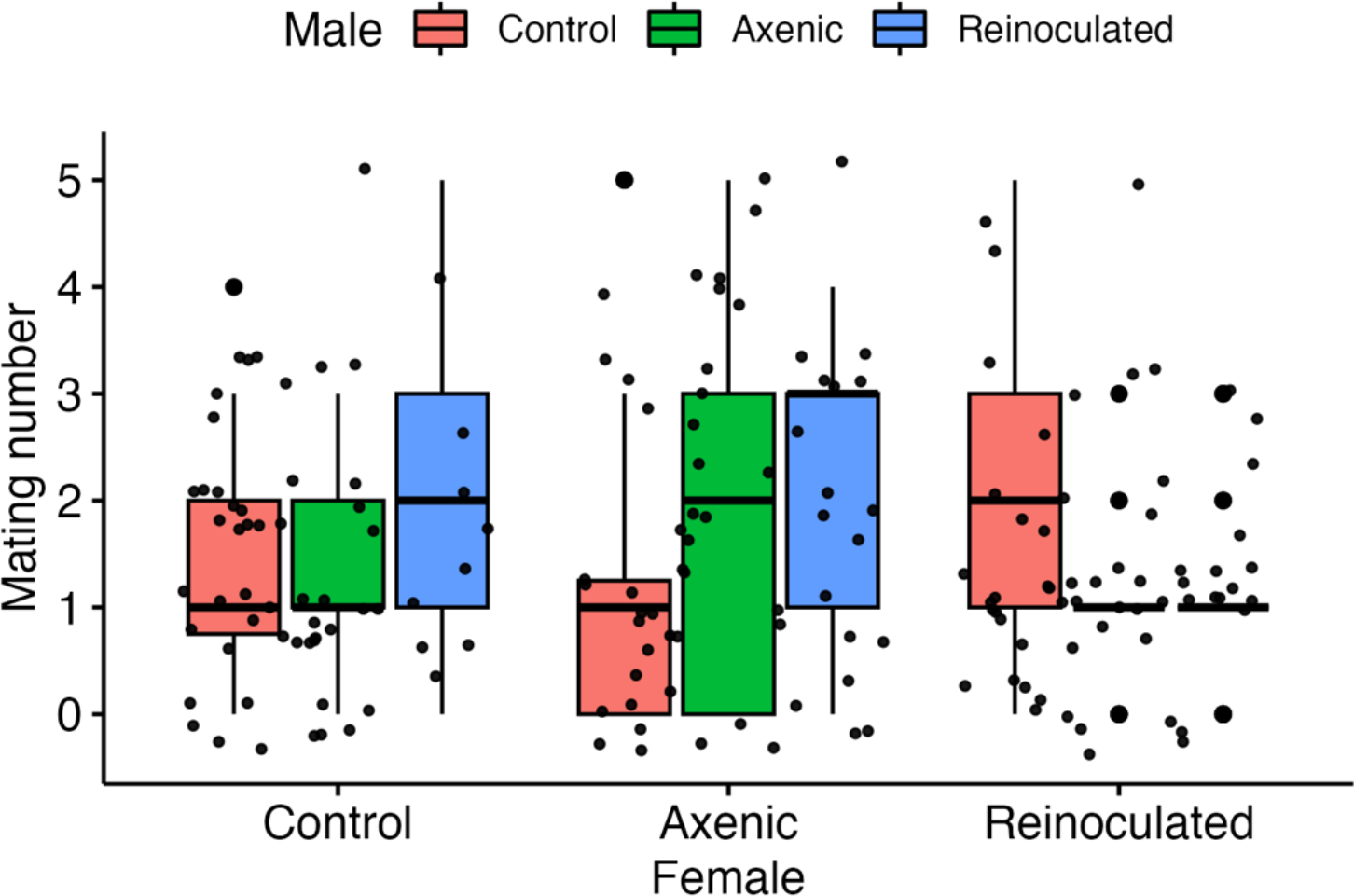
Number of matings for pairs between control, axenic, and reinoculated female and male hive beetles. Boxplots depict the median, the 25^th^ and 75^th^ percentile with the whiskers extended to the minimum and maximum data points.

### 3.4 Mating latency

The interaction between female and male treatment influenced the average mating duration (GLM (gaussian): Male: *F* = 5.695, *df* = 2, *p* < 0.01; Female: F =, *df* = 2, *p* < 0.01; Male*Female: *df* = 4, *p* = 0.043). For axenic females, the latency period to mate was the shortest when they mated with axenic or reinoculated males (Figure 3).

**Figure 3.**
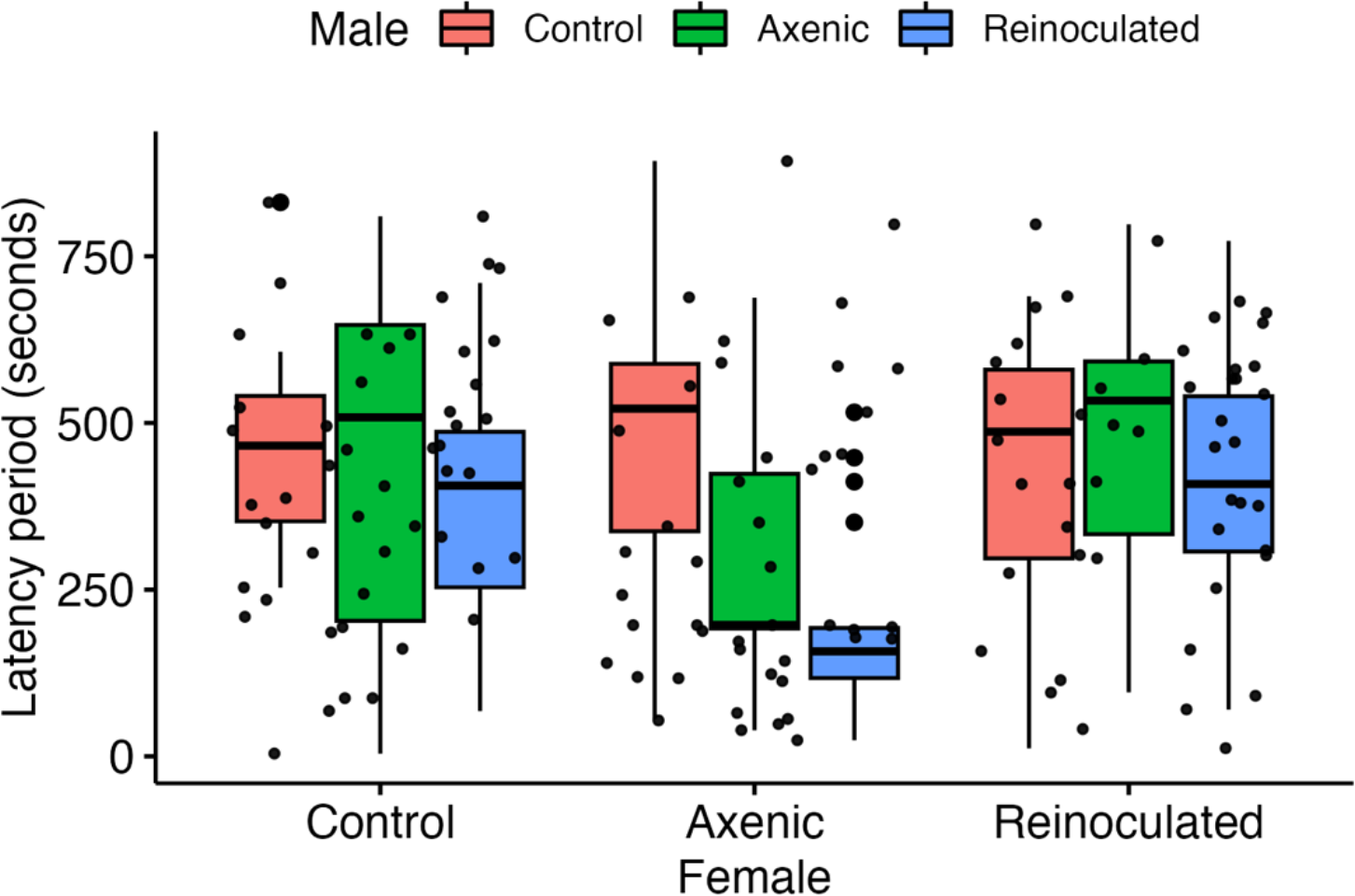
Latency period by pairs between control, axenic, and reinoculated female and male hive beetles. Boxplots depict the median, the 25^th^ and 75^th^ percentile with the whiskers extended to the minimum and maximum data points.

### 3.5 Mating duration

The interaction between sex and treatment influenced the average mating duration (GLM (gaussian): Male: *F* = 0.707, *df* = 2, *p* = 0.495; Female: *F* = 1.034, *df* = 2, *p* = 0.358; Male*Female: *F* = 2.876, *df* = 4, *p* < 0.05). The mating duration was the longest when control males mated with control females (Figure 4).

**Figure 4.**
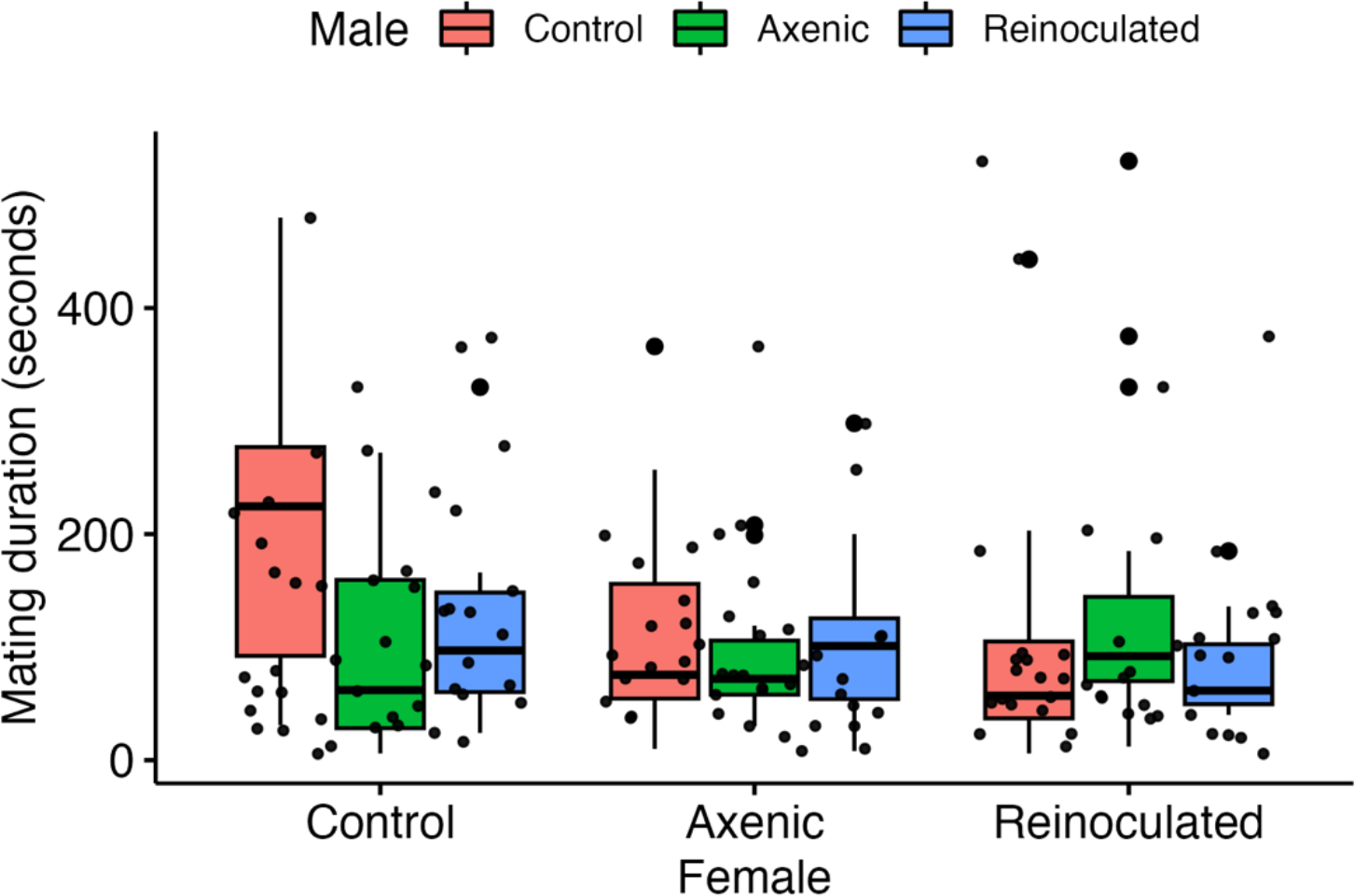
Mating duration for pairs between control, axenic, and reinoculated female and male hive beetles. Boxplots depict the median, the 25^th^ and 75^th^ percentile with the whiskers extended to the minimum and maximum data points.

## 4 DISCUSSION

In this study, we investigated the effects of manipulating the gut bacteria of male and female small hive beetles on mating behaviour and reproductive output. We found sex-specific effects of the gut bacteria manipulation on mating latency, mating number, mating duration, and offspring number with an adverse effect of female gut bacteria removal on larval production. The findings of this study show the intricate relationship between the insect and its gut bacteria, emphasizing the essential role of commensals on insect reproduction.

Manipulation of both male and female gut bacterial composition affected mating latency; however, no effect on mating duration was observed [but see Morimoto et al. (2017) where male gut bacteria community significantly impacted the duration of mating in *Drosophila melanogaster*]. We also observed that females with very few bacteria in their digestive tract produced less larvae than females with an intact or reinoculated gut bacteria. Similarly, previous work has demonstrated that females who lack a microbiome exhibit changes in their ability to produce offspring compared to females who possess a microbiome (Nguyen, Than, Dinh, Morimoto, & Ponton, 2020). Here we have not identified the bacterial strains involved in this decrease in fecundity, however, it has been shown that *D. melanogaster* females lacking *Acetobacter* species in their gut suffer a fecundity drop (Elgart et al., 2016; Morimoto et al., 2017). Further, Qiao, Keesey, Hansson, & Knaden (2019) have shown that fecundity requires a constant supply of *Saccharomyces cerevisiae* to be maintained. Female insects may initiate metabolic changes after mating to meet the heightened energy requirements necessary for egg production (Apger-McGlaughon & Wolfner, 2013; Cognigni, Bailey, & Miguel-Aliaga, 2011; McMillan et al., 2018; Reiff et al., 2015; Uchizono, Tabuki, Kawaguchi, Tanimura, & Itoh, 2017). A lack a gut bacteria might affect their capacity to regulate the metabolic shifts that are essential for sustaining egg production (Elgart et al. 2016). This may be a consequence of a change in foraging behaviour as it was observed in fruit flies (*Bactrocera tryoni*) when gut bacteria were manipulated (Nguyen, Dinh, Morimoto, & Ponton, 2021).

The microbiome of a female’s partner can impact the degree of reproductive success in *Drosophila* (Delbare, Ahmed-Braimah, Wolfner & Clark, 2020; Morimoto, Simpson & Ponton, 2017). Our study also revealed that the gut bacteria of male beetles played a role in influencing the fecundity of females. Larval production was lower in control females that mated with an axenic male. Previous research has shown that gnotobiotic *Drosophila* males, only carrying the bacterium *Lactobacillus plantarum* in their gut, have longer copulation duration and induce higher short-term egg-laying behaviour in their mates (Morimoto, Simpson & Ponton, 2017). When males only carried the bacterium *Acetobacter pomorum*, the ability of the mating pair to produce offspring was however impaired (Morimoto, Simpson & Ponton, 2017). These results indicate that the presence of particular strains of gut bacteria is crucial not only for female but also for male reproductive success (see also (Delbare, Ahmed-Braimah, Wolfner & Clark, 2020)). It is unclear in our study if the lack of the same symbiont affects similarly male and female. The composition of the gut bacteria can vary between sexes in insect species with some bacterial strains specifically found in the digestive tract of one sex but not the other (Chen et al., 2016; Tang, Adler, Vogel, & Ping, 2012). More investigation is required to identify which bacterial strains are specifically involved in small hive beetle, male and female, reproduction.

Based on our findings, manipulating the bacterial community associated with small hive beetles could be a potential strategy for controlling this pest. By identifying key bacteria involved in mating behaviour and reproduction, it may be possible to develop methods to disrupt or manipulate these processes, ultimately suppressing the small hive beetle populations. Manipulation of insect-associated microbiomes can be a vital tool for pest management (Qadri, Short, Gast, Hernandez, & Wong, 2020). For example, ingesting antibiotics such as tetracycline and penicillin has been demonstrated to induce sterility in tsetse flies by disrupting their essential mutualistic relationship with *Wigglesworthia* bacteria, and this disruption also hampers the growth of young ticks and reduces the reproductive capabilities of mature ticks due to a reduction in their symbiotic microbial population (Zhong, Jasinskas, & Barbour, 2007). Hence, this study represents an important initial step in unraveling the intricate relationship between gut bacteria and small hive beetle behaviour, paving the way for future investigations and the development of innovative pest management strategies. Finally, due to the intimate relationship between small hive beetles and bees, understanding the physiological roles of beetle gut bacteria might inform on the relationship between bees and their symbiotic bacteria.

## AUTHOR CONTRIBUTIONS

**Md Jamil Hossain Biswas**: Conceptualization (lead); funding acquisition (lead); methodology (lead); conducted the experiment (lead); data curation (lead); formal analysis (equal); visualization (lead); writing – original draft (lead); writing – review and editing (equal). **Md Kawsar Khan**: Conceptualization (supporting); methodology (supporting); conducted the experiment (equal); data curation (supporting); writing – review and editing (equal). **Sahareh Bashiribod**: Methodology (supporting); conducted the experiment (equal); writing – review and editing (equal). **Fleur Ponton**: Conceptualization (lead); funding acquisition (supporting); methodology (lead); conducted the experiment (supporting); data curation (supporting); formal analysis (lead); writing – review and editing (equal).

## ACKNOWLEDGMENTS

This research project was funded by Md Jamil Hossain Biswas’ COVID Recovery Fellowship. We acknowledge Mr. The Anh Than and Ms. Kieu Diem Quynh Nguyen for their help in experiments.

## DATA AVAILABILITY STATEMENT

Upon acceptance of this manuscript, the data will be available in ‘figshare’.

## ETHICS STATEMENT

None required

